# Formate from THF-C1 metabolism induces the AOX1 promoter in formate dehydrogenase-deficient *Pichia pastoris*

**DOI:** 10.1101/2024.05.29.596193

**Authors:** Cristina Bustos, Julio Berrios, Patrick Fickers

**Affiliations:** Microbial Processes and Interactions, TERRA Teaching and Research Centre, Gembloux Agro-Bio Tech, University of Liege, Gembloux, Belgium; School of Biochemical Engineering, Pontificia Universidad Católica de Valparaíso, Av Brasil 2085, Valparaiso 2340000, Chile

## Abstract

In *Pichia pastoris* (*Komagataella phaffii*), formate is a recognized alternative inducer to methanol for expression systems based on the AOX1 promoter (pAOX1). By disrupting the formate dehydrogenase encoding *FDH1* gene, we converted such a system into a self-induced one, as adding any inducer in the culture medium is no longer requested for pAOX1 induction. In cells, formate is generated from serine through the THF-C1 metabolism, and it cannot be converted into carbon dioxide in an *fdh1*Δ strain. Under non-repressive culture conditions, such as on sorbitol, the intracellular formate generated from the THF-C1 metabolism is sufficient to induce pAOX1 and initiate protein synthesis. This was evidenced for two model proteins, namely intracellular eGFP and secreted CalB lipase from *C. antarctica*. Similar protein productivities were obtained for an *fdh1*Δ strain on sorbitol and a non-disrupted strain on sorbitol-methanol. Considering a *P. pastoris fdh1Δ* strain as a workhorse for recombinant protein synthesis paves the way for the further development of methanol-free processes in *P. pastoris*.

## Introduction

The methylotrophic yeast *Pichia pastoris* (*Komagataella phaffii*) is a well-established and reliable cell factory for producing recombinant proteins (rProt) (Ergün *et al*., 2021; Barone *et al*., 2023). The expression systems used typically and historically rely on the regulated promoter from the alcohol oxidase 1 gene (pAOX1). This promoter is repressed during cell growth on glycerol, while pAOX1 induction and thus, rProt synthesis, is triggered by adding an inducer to the culture medium, typically methanol (Ergün *et al*., 2021; Bustos *et al*., 2022). Although widely used, including on an industrial scale, methanol presents several technical challenges that are difficult to overcome in practice. It is highly flammable and can become toxic to cells at high concentrations due to the accumulation of toxic methanol catabolic products such as formaldehyde (Berrios *et al*., 2022) (Fig. S1). Moreover, its oxidation by alcohol oxidases in peroxisomes requires oxygen, thereby increasing the cellular oxygen demand compared to other carbon sources. Additionally, methanol catabolism generates heat, which must be dissipated, thereby increasing the operational costs, especially for large-scale production processes (Krainer *et al*., 2012; Niu *et al*., 2013).

Formate, an intermediate metabolite of the methanol dissimilation pathway (Hartner & Glieder, 2006, Fig. S1), has emerged as an interesting alternative inducer to methanol for rProt synthesis in *P. pastoris*. It is produced from formaldehyde by formaldehyde dehydrogenase (Fld) before being converted into carbon dioxide by formate dehydrogenase (Fdh). Compared to methanol, formate is a more sustainable inducer that can be efficiently produced through the electrochemical conversion of carbon dioxide (Jhong *et al*., 2013; Cotton *et al*., 2020). The ability of formate to induce pAOX1 has been demonstrated (Tyurin and Kozlov, 2015; Jayachandran *et al*., 2017; Singh and Narang, 2020). However, one of the primary limitations of formate is its poor ability to be catabolized by *P. pastoris*. To address this constraint, an engineering strategy has been developed through the co-overexpression of genes encoding *Escherichia coli* acetyl-CoA synthase, *Listeria innocua* acetaldehyde dehydrogenase, and the transcription factor Mit1. This engineering effort led to an increase in rProt production (i.e. xylanase, Liu et al., 2022).

In cells, formate is also an intermediate of the tetrahydrofolate-mediated one-carbon (THF-C1) metabolism involved in several anabolic pathways, including the *de novo* synthesis of purines (Fig. 1, Kastanos et al., 1997; Piper et al., 2000). It is obtained from cytoplasmic serine by the action of the serine hydroxymethyltransferase (Shm) and the trifunctional C1-tetrahydrofolate synthase (Mis1) (Fig. 1). In the THF-C1 metabolism, formate serves as a shuttle for C1 units between the cytoplasm and the mitochondrion, as THF derivatives cannot cross the mitochondrial membrane (Kastanos *et al*., 1997).

**Figure 1:**
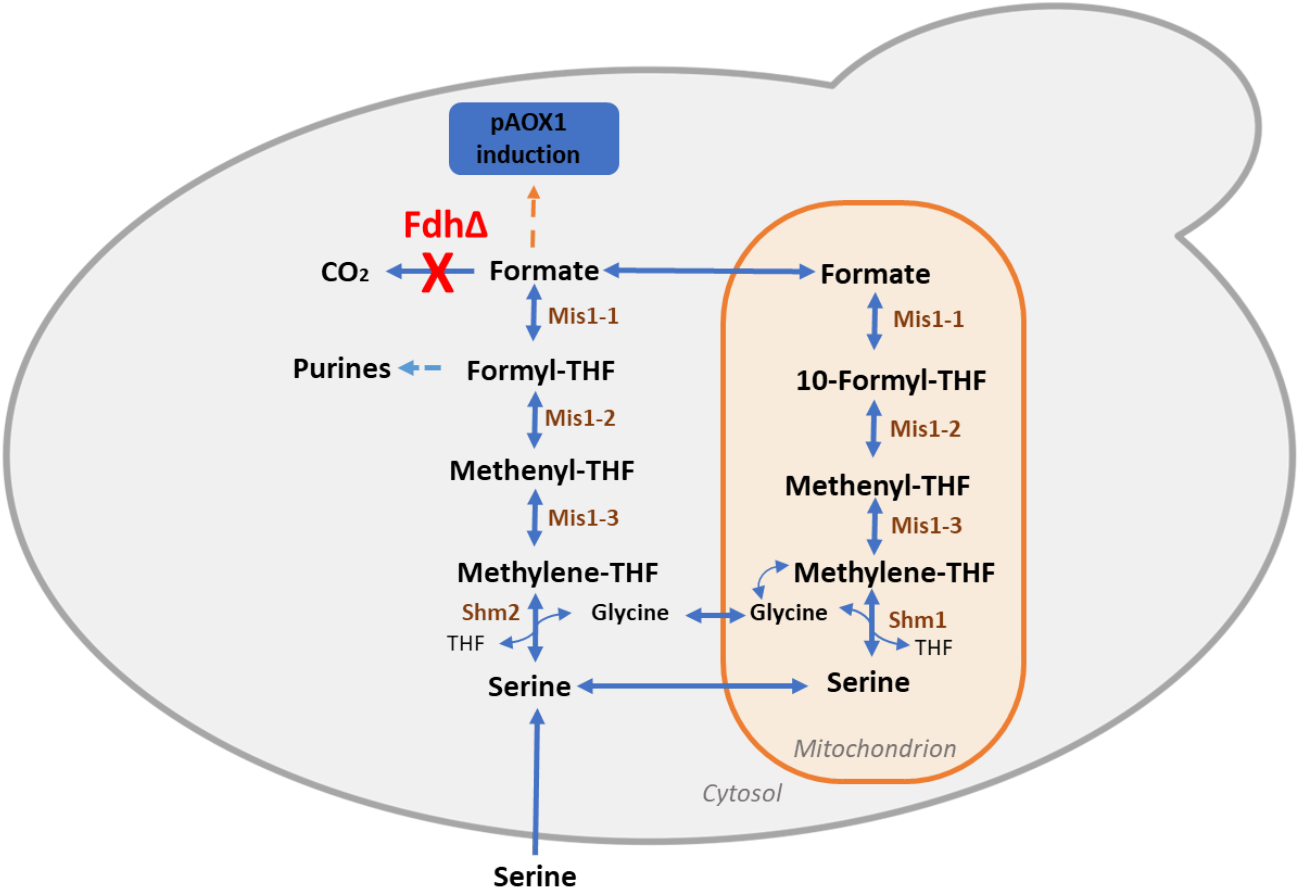
Tetrahydrofolate (THF) mediated one-carbon (THF-C1) metabolism in yeast. Fdh, formate dehydrogenase; Shm, serine hydroxymethyltransferase; Mis1, trifunctional C1-tertahydrofolate synthase (Mis1-1, Mis1-2, Mis1-3); THF, tetrahydrofolate. FDH gene knockout is mentioned in red, and enzymes involved in pathways are shown in brown.

Herein, we aim to investigate the regulation of pAOX1 by formate in an *FDH1* knockout (*fdh1*Δ) strain. From our investigation, it became evident that endogenous formate from THF-C1 metabolism was sufficient to trigger pAOX1 induction in an *fdh1Δ* mutant grown under non-repressive culture conditions (i.e. in the presence of sorbitol) without any supplementation of inducer. In those conditions, any pAOX1-based expression system could be potentially converted into a self-induced one.

### Experimental procedures

#### Strains and media and culture conditions

The *P. pastoris* and *Escherichia coli* strains used are listed in Table 1 and S1, respectively. *E. coli* was cultivated at 37°C in Luria-Bertani medium (LB), supplemented with antibiotics as follows: 100 µg ml^-1^ ampicillin, 50 µg ml^-1^ kanamycin, 25 µg ml^-1^ zeocin, or 50 µg ml^-1^ hygromycin. *P. pastoris* strains were cultivated at 30°C either in YPD medium (containing 20 g l^-1^ glucose, 10 g l^-1^ Difco yeast extract, and 10 g l^-1^ Difco bacto peptone) or YNB medium (containing 1.7 g L^-1^ Difco YNB w/o ammonium chloride and amino acids, 5 g l^-1^ NH4Cl and, 0.4 mg l^-1^ biotin, 100mM potassium phosphate buffer, pH 6.0) supplemented with as follows: 10 g l^-1^ sorbitol and 2 g l^-1^ Difco casamino acid (YNBSC), 10 g l^-1^ sorbitol, 5.1 g l^-1^ methanol and 2 g l^-1^ Difco casamino acid (YNBSMC); 10 g l^-1^ sorbitol, 10.8 g l^1^ formate and2 g l^-1^ Difco casamino acid (YNBSF), 10 g. l^-1^ sorbitol (YNBS), 10 g l^-1^ sorbitol and 5.1 g l^-1^ methanol (YNBSM); 10 g l^-1^ sorbitol and 10.8 g l^1^ formate (YNBSF); 10 g l^-1^ glycerol (YNBG); 6.3 g l^-1^ methanol and 4.0 g l^-1^ sorbitol (YNBMS); 10 g l^-1^ sorbitol and 2 g l^-1^ serine (YNBSS), 10 g l^-1^ sorbitol with 2 g l^-1^ glycine (YNBSG). *P. pastoris* transformants were selected on YPD agar plates, supplemented with antibiotics as follows when requested: 25 µg ml^1^ zeocin (YPD-Zeo), 200 µg ml^-1^ hygromycin (YPD-Hygro), 500 µg ml^1^ geneticin (YPD-Genet) or 100 µg ml^1^ nourseothricin (YPD-Nat).

**Table 1.** *Pichia pastoris* strains used in this study.

| Name | Parental strain, Genotype | Source/Reference |
| --- | --- | --- |
| RIY232 | GS115, *HIS4* | Theron et al (2019) |
| RIY230 | GS115, *pAOX1-eGFP* | Velastegui et al (2019) |
| RIY308 | GS115, *pAOX1-αMF-CalB* | Velastegui et al (2019) |
| RIY536 | RIY230, *fdh1Δ*, *pAOX1-eGFP*, Zeo+ | This work |
| RIY537 | RIY308, *fdh1Δ*, *pAOX1-αMF-CalB*, Zeo+ | This work |
| RIY540 | RIY536, *fdh1Δ*, *pAOX1-eGFP* | This work |
| RIY561 | RIY537, *fdh1Δ*, *pAOX1-αMF-CalB* | This work |
| RIY624 | RIY540, *fdh1Δ*, *pAOX1-eGFP*, *pGAP-FDH1*, Zeo+ | This work |
| RIY641 | RIY540, *fdh1Δ*, *shm1Δ*, *pAOX1-eGFP*, Nat+ | This work |
| RIY640 | RIY540, *fdh1Δ*, *shm2Δ*, *pAOX1-eGFP*, Zeo+ | This work |
| RIY642 | RIY540, *fdh1Δ*, *shm1Δ*, *shm2Δ*, *pAOX1-eGFP*, Zeo+, Nat+ | This work |

For all cultures, a first preculture inoculated from a single colony was performed for 12 h at 30°C and 150 rpm in a 250 ml shake flask containing 25 ml of liquid YPD medium. After centrifugation at 9000 g for 5 min, the cells were washed with phosphate-buffered saline (0.1 M, pH 6) before being used to inoculate a second preculture in the same conditions in YNB media supplemented as described above. Cultures were performed in 24-square deep well plates (System Duetz, Enzyscreen) as described elsewhere containing 1.5 ml of medium (Sassi et al. 2016), in 50 ml shake flasks (5 ml medium) or in microbioreactor (BioLector 2, m2p-labs, Baesweiler, Germany). For that purpose, 48-well Flower plates (M2P-MTP-48-B, Beckman Coulter Life Sciences, USA) containing 1 ml of medium were used. Cultures were operated for 60h with a relative humidity of 85%, under constant agitation at 1000 rpm. Every 10 minutes, biomass was monitored using scattered light intensity at a wavelength of 620 nm while cell fluorescence was quantified at 520 nm (excitation at 488 nm). The gain was set as 2 for biomass and 4 for fluorescence. Specific fluorescence was obtained by dividing the fluorescence value by the biomass value. It was expressed in specific fluorescence units (sFU). All cultures were seeded at an initial optical density at 600 nm of 0.5 from cells grown in the second preculture. Cultures in 24-square deep well plates were performed with three biological replicates, whereas cultures in the BioLector were performed with two biological replicates, each supported by two technical replicates, resulting in a total of four replicates.

#### General genetic techniques

Standard media and techniques were used for *E. coli* (Sambrook and Russell, 2001). Restriction enzymes, DNA polymerases, and T4 DNA ligase were obtained from New England Biolabs (NEB, Ipswich, MA, USA) or Thermo Scientific (Thermo Scientific, Waltham, MA USA). Primers for PCR and qPCR were synthesized by Eurogentec (Seraing, Belgium, Table S2). Vector TopoBluntII and pGEMTeasy were from Invitrogen (Waltham, Massachusetts, United States) and Promega (Madison, Wisconsin, United States), respectively. Genomic DNA was purified using a Genomic DNA Purification kit (Thermo Scientific, Waltham, MA USA). DNA fragments were purified from agarose gels using a NucleoSpin Gel and a PCR clean-up kit (Machery-Nagel, Düren, Germany). DNA sequencing was performed by Eurofin Genomic (Eurofin, Ebersberg, Germany). Quantitative PCR (qPCR) were performed as described elsewhere with primers listed in Table S1, using the actin gene as a reference. Total RNA was extracted using the NucleoSpin RNA Plus kit (Machery-Nagel, Düren, Germany). qPCR was performed using the Luna Universal qPCR Master Mix and the Step OnePlus Real-Time PCR system (Thermo Scientific, Waltham, MA, USA). Primers and plasmid designs were performed using the software Snapgene (Dotmatics, USA). Vectors were constructed using the GoldenPiCS Kit (Prielhofer et al., 2017, Addgene kit #1000000133). *P. pastoris* was transformed as described by Lin-Cereghino et al., (2005).

#### Construction of plasmids and P. pastoris strains

To construct the gene disruption cassettes, a ∼ 1kb fragment upstream of the start codon (Pro-gene) and ∼ 1kb fragment downstream of the stop codon (Term-gene) of the genes PAS_chr3_0932 (*FDH1*), PAS_chr4_0587 (*SHM1*) and PAS_chr4_0415 (*SHM2*) were PCR-amplified using *P. pastoris* GS115 genomic DNA as a template. The primer pairs used to amplify Pro-gene and Term-gene were P.fdh1-Fw/P.fdh1-Rv and T.fdh1-Fw/T.fdh1-Rv for *FDH1*, P.shm1-Fw/P.shm1-Rv and T.smh1-Fw/T.shm1-Rv for SHM1; and P.shm2-Fw/P.shm2-Rv and T.smh2-Fw/T.shm2-Rv for *SHM2*. The zeocin and nourseothricin selection markers were amplified from plasmids D12-BB3aZ_14 and E6-BB3aN_14 (Table S1), used as a template with primer pairs BleoR.fdh1-Fw/BleoR.fdh1-Rv, Nat.shm1-Fw/Nat.shm1-Rv and BleoR.shm2-Fw/ BleoR.shm2-Rv and subsequently used to construct the *FDH1, SHM1* and *SHM2* disruption cassettes, respectively. The *FDH1* disruption cassette (P_fdh1-Bleo.R-T_fdh1) was obtained by Golden Gate assembly using BsaI as restriction enzyme. The *SHM1* and *SHM2* disruption cassettes (Pro_gene-Selection Marker-Term_gene) were obtained by an overlapping PCR using the corresponding purified Pro_gene, selection marker, Term_gene fragment as templates and primer pairs P.fdh1-Fw/T.fdh1-Rv, P.shm1-Fw/T.shm1-Rv and P.shm2-Fw/T.shm2-Rv, respectively. The resulting ∼ 3.2 kb fragments were cloned into the pGEMT-Easy vector or Blunt II-Topo vector to generate plasmids RIP 369 (*FDH*), RIP491 (*SHM2*), RIP492 (*SHM1*) (Table S1). The FDH1 disruption cassette from plasmid RIP369 was subsequently used to transform strains RIY230 (pAOX1-eGFP) and RIY308 (pAOX1-αMF-CalB) to yield strains RIY536 (*fdhΔ*, pAOX1-eGFP, Zeo+) and RIY537 (*fdhΔ*, pAOX1-αMF-CalB, Zeo+), respectively. Construction of strains RIY230 and RIY308 were described in Velastegui et al., (2019). The SMH1 and SHM2 disruption cassettes from plasmids RIP492 and RIP491 were used to transform strain RIY540 to generate the strains RIY639, RIY641 and RIY642 (Table 1). The disruption cassettes were released from the corresponding plasmid by SacI restriction. Transformants were selected on YPD-Zeo and YPD-Nat, according to the corresponding marker. For marker rescue, strains RIY536, and RIY237 were transformed with the replicative vector RIP396 (pKTAC-Cre) and transformants were selected YPD-Genet. The resulting strains were RIY540 (*fdhΔ*, pAOX1-eGFP) and RIY561 (*fdhΔ*, pAOX1-αMF-CalB). To construct the *FDH1* expression vector, the GoldenPiCS system was used (Prielhofer *et al*., 2017). Internal BpiI recognition sequence in gene PAS_chr3_0932 (*FDH1*) was removed by overlapping PCR using pairs Fdh1-Fw/Fdh1.BpiI-Rv and Fdh1.BpiI-Fw/Fdh1-Rv using *P. pastoris GS115* genomic DNA as a template. The resulting PCR product was cloned into plasmid A2 (BB1-23) at BsaI restriction site to yield plasmid RIP465. Plasmid RIP466 (pGAP-FDH1-ScCYC1tt) was constructed by Golden Gate assembly from the plasmids RIP465, A4 (BB1_12_pGAP), C1 (BB1_34_ScCYC1tt) and E1. (BB3eH_14) using BpiI as the restriction enzyme. After PmeI digestion and purification, plasmid RIP466 was used to transform the RIY540 strain (*fdhΔ*, pAOX1-eGFP) to yield the RIY624 strain (pAOX1-EGFP, pGAP-FDH). Transformants were selected on YPD-Hygro. Correctness of the disruption mutant genotype was confirmed by analytical PCR on the genomic DNA of the different disrupted strains. For gene disruption, the forward primers annealed upstream of the Pro-genes, namely Up.fdh1-Fw, Up.shm1-Fw, Up.shm2-Fw, for genes *FDH1, SHM1* and *SHM2*, respectively, while the reverse primers annealed within the selection marker, namely BleoR.Int-Rv for genes *FDH1* and *SHM2*, and Nat.shm1-Rv for gene *SHM1*. As further confirmation, forward primers that annealed within the selection marker BleoR.Int-Fw for gene *FDH1* and *SHM2, and* Nat.Int-Fw for gene *SHM1* and reverse primers that annealed downstream of the Term-gene, namely Dw. fdh1-Rv, Dw.shm1-Rv, Dw.shm2-Rv, for gene *FDH1, SHM1* and *SHM2*, respectively, were used. To confirm the excision of the selection marker in the *fdh1*Δ strain RIY536, primer pairs P.fdh1-Fw/T.fdh1-Rv were used. To verify the genotype of the RIY624 strain (*FDH1* complemented strain), primers pGAp.Int-Fw and Cyc1t.Int-Rv that annealed in the pGAP and the ScCYC1tt region were used.

#### Analytical methods

Cell growth was monitored either by optical density at 600nm (OD_600_) or dry cell weight (DCW) as previously described (Carly *et al*., 2016). Methanol, sorbitol, and glycerol concentrations were determined by HPLC (Agilent 1100 series equipped RID detector, Agilent Technologies, Santa Clara, CA, USA) using an Aminex HPX-87H ion-exclusion column (300 × 7.8 mm Bio-Rad, Hercules, CA, USA). Compounds were eluted from the column at 65 °C with a flow rate of 0.5 ml min^−1^ and using a 5 mM H_2_SO_4_ solution as the mobile phase.

Intracellular eGFP fluorescence was quantified using a BD Accuri C6 Flow Cytometer (BD Biosciences, San Jose, CA, USA) as described elsewhere(Sassi *et al*., 2016). For each sample, 20,000 cells were analyzed using the FL1-A and FSC channels, and FL1-A/FSC dot plots were analyzed using the CFlowPlus software (Accuri, BD Biosciences). A threshold of 5800 fluorescence units (FU) on FL1-A channel was applied to eliminate the noise for endogenous fluorescence from the cells. To calculate the total value of fluorescence in the cell population, the FL1-A median value (i.e., the eGFP fluorescence) was multiplied by the fraction of cells with eGFP fluorescence (i.e., induced cells). It was expressed in total fluorescence unit (TFU). Spectrophotometric analysis of eGFP was performed on SpectraMax M2 (Molecular Devises, San Jose San Jose, CA, USA) using λex and λem at 488 and 535 nm, respectively. Measurements were taken after 30 s of sample shaking. Signal gain was set to 225, and the number of light flashes was set to 30. Specific eGFP fluorescence was expressed as specific fluorescence units (SFU), i.e., as fluorescence value normalized to biomass related to optical density at 600 nm (OD600) of 0.5.

The lipase activity in the culture supernatant was determined by monitoring the hydrolysis of p-nitrophenylbutyrate (p-NPB) as described elsewhere (Fickers *et al*., 2003). The release of para-nitrophenol was monitored at 405 nm using a SpectraMax M2 (Molecular Devices, San Jose, CA, USA). All lipase activity assays were performed at least in triplicate. One unit of lipase activity was defined as the amount of enzyme releasing 1 µmol p-nitrophenol per minute at 25 °C and pH 7.2 (εPNP = 0.0148 µM^−1^.cm^−1^).

#### Fluorescence microscopy

Microscopy was performed with a Nikon Eclipse Ti2-E inverted automated epifluorescence microscope (Nikon Eclipse Ti2-E, Nikon France, France) equipped with a DS-Qi2 camera (Nikon camera DSQi2, Nikon France, France), a 100× oil objective (CFI P-Apo DM Lambda 100× Oil (Ph3), Nikon France, France). The GFP-3035D cube (excitation filter: 472/30 nm, dichroic mirror: 495 nm, emission filter: 520/35 nm, Nikon France, Nikon) was used to visualize eGFP. Prior observation, cells were washed with phosphate buffer saline and diluted at a cell concentration of 0.5 gDCW l^-1^. For image processing, ImageJ software was used (Collins, 2007; Schneider *et al*., 2012).

## Results and discussion

### AOX1 promoter activity is upregulated by formate

In methylotrophic yeasts, formate is an intermediate of the methanol dissimilation pathway (Hartner & Glieder, 2006, Fig. S1). Studies have demonstrated the efficacy of formate as both an inducer and a carbon source to produce recombinant proteins (rProt) in *P. pastoris* (Singh and Narang, 2020; Liu, Li, *et al*., 2022; Liu, Zhao, *et al*., 2022). Herein, an enhanced green fluorescent protein (eGFP) reporter system was used to probe the regulation of the *AOX1* gene promoter (p*AOX1*) by formate. For this purpose, the RIY230 strain (*pAOX1-eGFP*, hereafter *FDH1* strain, (Velastegui *et al*., 2019) was grown on sorbitol (YNBS) supplemented or not with methanol or formate (YNBSC, YNBSMC, and YNBSFC, respectively). Sorbitol was selected as the carbon source since it is known as non-repressive for pAOX1 (Niu *et al*., 2013). Specific eGFP fluorescence was monitored in cells by flow cytometry at the end of the growth phase (i.e., 18h) and during the stationary phase (i.e., 24h). As shown in Fig. 2, the pAOX1 induction levels (eGFP signal) were low on sorbitol medium (16601 and 9340 TFU, respectively). They were remarkedly higher and in the same range for cells grown on sorbitol-methanol (61207 and 48518 TFU, respectively) or sorbitol-formate (50488 and 52960 TFU, respectively). These observations contrast with a recent report on similar experiments conducted on glycerol-based defined media (YNBG), where eGFP-specific fluorescence levels were reported as 4.1-fold lower in the presence of formate compared to methanol (Feng *et al*., 2022). This demonstrates that formate can substitute methanol for pAOX1 induction, at least in a sorbitol-minimal medium.

**Figure 2:**
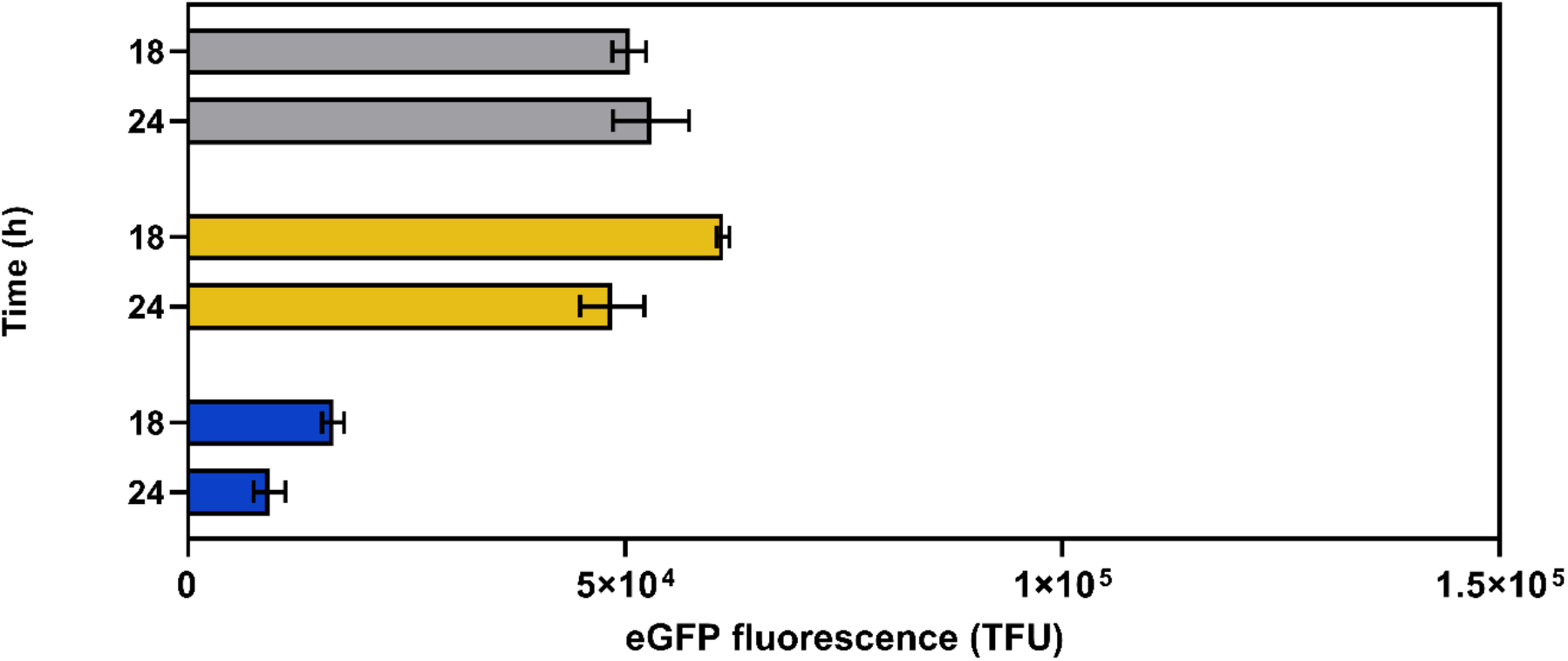
eGFP fluorescence of RIY230 strain (pAOX1-eGFP, FDH1 strain) after 18h and 24 h of growth in YNB minimal medium containing sorbitol and formate (YNBSFC, grey), sorbitol and methanol (YNBSMC, yellow), and sorbitol (YNBSC, blue). Fluorescence was quantified by flow cytometry on 20,000 cells and expressed as TFU (total fluorescence, see materials and method for calculation details). Data are the mean and standard deviation of triplicate cultures conducted in deep well plates.

### Formate can be used as a free inducer in an fdh1Δ strain

In the methanol dissimilation pathway, formate is converted to carbon dioxide by formate dehydrogenase (Fdh, Fig. S1). In methylotrophic yeasts, including *P. pastoris*, Fdh was shown as non-essential for cell survival. However, the growth of a formate dehydrogenase knockout mutant (*fdh1*Δ) is remarkedly reduced in a methanol-based medium (Guo *et al*., 2021). Moreover, an *fdh1*Δ strain exhibits a heightened sensitivity to the accumulation of formate in the medium, indicating that the primary physiological function of Fdh is more related to the detoxification of intracellular formate rather than energy generation (Sibirny *et al*., 1990; Sakai *et al*., 1998). Therefore, the knockout of gene *FDH1* in *P. pastoris* would render formate a free pAOX1 inducer in non-repressive conditions (in the presence of sorbitol). For that purpose, the *FDH1* knockout RIY540 strain (*fdh1Δ, pAOX1-eGFP*, hereafter *fdh1*Δ, Table 1) was constructed. It was grown on sorbitol, sorbitol-methanol, or sorbitol-formate (YNBSC, YNBSMC, and YNBSFC, respectively), and the eGFP fluorescence was quantified by flow cytometry after 18 h and 24 h of culture. On sorbitol-methanol (YNBMC), eGFP fluorescence signals were on average for both sampling times in the same range for the *fdh1Δ* and *FDH1* strains (i.e. 42715 and 54862 TFU, respectively; Fig. 2 and 3). This demonstrates that the knockout of *FDH1* has no impact on the strength of the pAOX1 induction level by methanol. By contrast, on sorbitol-formate, the fluorescence signals were, on average, for both sampling times 1.9-fold higher for the *fdh1Δ* strain compared to the *FDH1* strain (i.e. 100359 and 51724 TFU, respectively). Therefore, preventing *P. pastoris* from dissipating formate into carbon dioxide yielded higher induction levels of pAOX1 on formate than on methanol. More importantly, for the *FDH1*-knockout strain, eGFP fluorescence signals were in the same range on sorbitol and sorbitol-formate on average for the two sampling times (i.e. 99632 and 100359 TFU, respectively). It was also 7.6-fold higher on average for the *fdh1Δ* strain compared to the *FDH1* strain on sorbitol (i.e., in the absence of any inducer; 99632 and 12970 TFU). These results were corroborated by quantifying the eGFP gene expression level for the *fdh1Δ* strain grown on sorbitol. It was 2.9 and 3.2-fold increased the *fdh1Δ* strain compared to the *FDH1* strain after 18 h and 24 h of growth, respectively (Fig. S2). Fluorescence microscopy also clearly showed a higher eGFP level for the knockout strain on sorbitol (i.e. without the addition of formate; Fig. S3).

**Figure 3.**
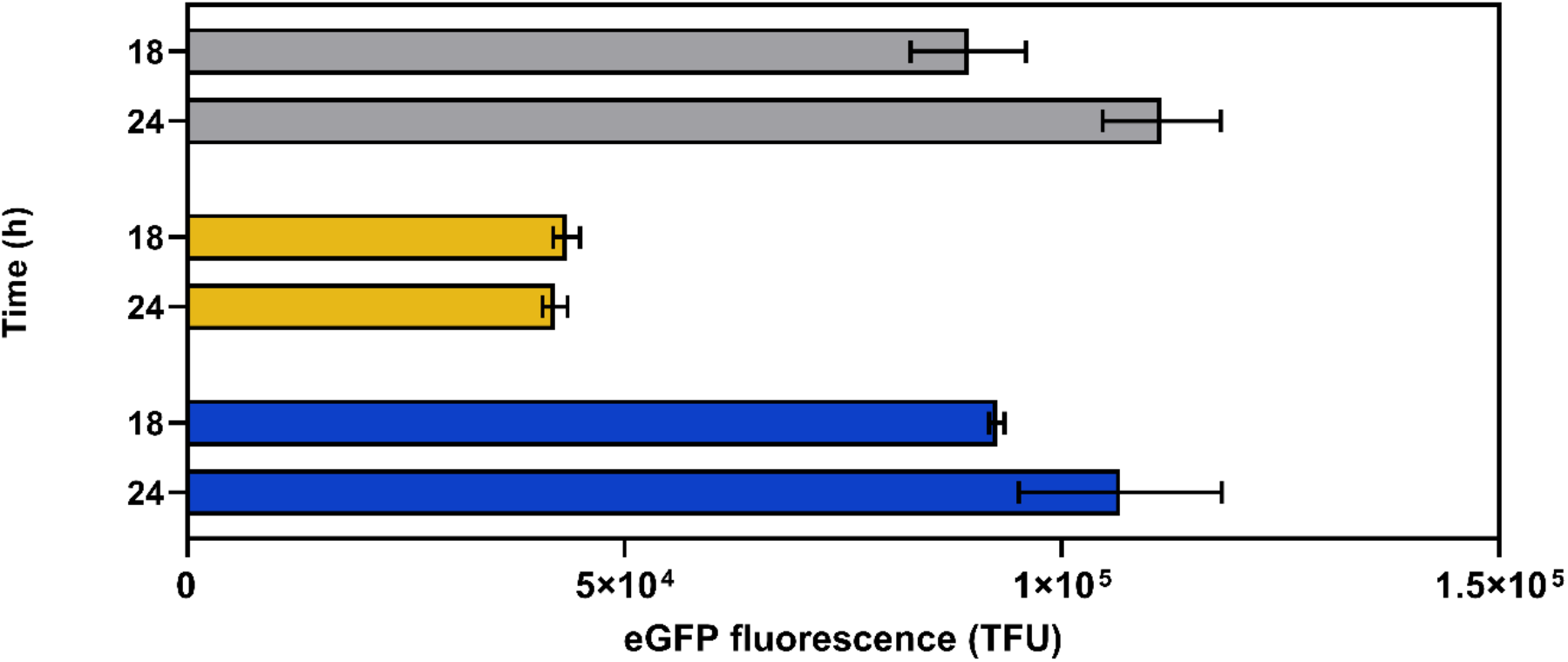
eGFP fluorescence of RIY540 strain (fdhΔ, pAOX1-eGFP, fdh1Δ strain) after 18h and 24 h of growth in YNB minimal medium containing sorbitol and formate (YNBSFC, grey), sorbitol and methanol (YNBSMC, yellow), and sorbitol (YNBSC, blue). Fluorescence was quantified by flow cytometry on 20,000 cells and expressed as TFU (total fluorescence, see materials and method for calculation details). Data are the mean and standard deviation of triplicate cultures conducted in deepwell plates.

### Complementation of the fdh1Δ strain restored the wild-type phenotype

To confirm that the phenotype of the *fdh1Δ* strain is related to the disruption of the gene PAS_chr3_0932, it was expressed under the control of the constitutive pGAP promoter in the RIY540 strain. The resulting RIY624 strain (*fdh1*Δ, *pAOX1-eGFP, pGAP-FDH*, hereafter *fdh1*Δ*-FDH1*) was grown on sorbitol (YNBS) together with FDH1 and *fdh1*Δ strains, used as negative and positive controls, respectively. The fluorescence level of the *fdh1*Δ*-FDH1* strain was reduced by 24-fold on average on two sampling times (18 h and 24 h) as compared to the *fdh1*Δ strain (i.e.87218 and 3657 TFU, respectively; Fig. S4). This demonstrates that the disruption of the *FDH1* gene is related to the phenotype of the knockout strain.

### Unravelling the origins of formate in a methanol-free environment

In the *fdh1*Δ strain, a strong increase in the pAOX1 induction level was observed under non-repressive culture conditions and in the absence of formate compared to the FDH1 strain (on sorbitol medium, YNBSC). This suggests that formate is generated in an alternative metabolic pathway and somehow accumulates intracellularly in the *fdh1*Δ strain. Besides the methanol dissimilation pathway, formate is generated from cytoplasmic serine in the THF-C1 metabolism by Shm1 and Mis1 enzymes (Fig. 1, Kastanos et al., 1997). In a *P. pastoris* wild-type strain, formate generated through that metabolism can, therefore, be consumed either by Mis1 to form formyl-THF or by Fdh to form carbon dioxide. As the disruption of gene *FDH1* prevent this conversion into carbon dioxide, formate may somehow accumulate intracellularly in the *fdh1*Δ strain, explaining thus the induction level of pAOX1 in non-repressive conditions. To verify this hypothesis, the expression of gene *FDH1* (as well as *FGH1* and *FLD*) was first confirmed by qPCR in cells grown on sorbitol (YNBS, Fig. S5). We then try to increase the intracellular formate formation through the THF-C1 pathway indirectly by the addition of serine in the culture medium. Therefore, FDH1 and *fdh1*Δ strains were grown in sorbitol-based media supplemented or not with serine (YNBS and YNBSS, respectively), and the specific fluorescence (i.e. normalized to biomass) was monitored over 60 h. For the FDH1 strain, the fluorescence signal remained at a constant and low level, similar to the RIY232 strain (GS115 prototroph), on both media and throughout the entire cultivation period (Fig. 4A & B). This suggests that pAOX1 is most probably not induced in those conditions in the FDH1 strain. By contrast, the fluorescence signal and thus pAOX1 induction level were remarkedly higher for the *fdh1*Δ strain, especially on a medium supplemented with serine. The specific fluorescence values for the *fdh1*Δ strain after 60 h of growth were 4.0 and 6.1-fold increased on sorbitol and sorbitol-serine, respectively, compared to the *FDH1* strain. Moreover, the addition of serine in the medium yielded for the *fdh1*Δ strain a 1.5-fold increased fluorescence signal compared to the non-supplemented medium. Similarly, we tried to decrease the intracellular formate formation through the THF-C1 pathway by growing the cell in the presence of glycine, as it has been reported as a Shm inhibitor (Piper *et al*., 2000). As shown in Fig 4C, the addition of glycine impaired pAOX1 induction for both strains for over 50 h. Gene PAS_chr4_0415 (*SHM2*) encoding cytoplasmic Shm was also disrupted in the *fdh1*Δ strain. The resulting RIY640 strain (*fdh1Δ, shm2Δ, pAOX1*-*eGFP*, hereafter *fdh1Δ*-*shm2Δ*) was grown on sorbitol in the presence or not of serine or glycine (YNBS, YNBSS and YNBSG, respectively). In all tested media, the specific fluorescence signal was markedly lower for *fdh1Δ*-*shm2Δ* strain as compared to the *fdh1Δ* strain (Fig. 4). By contrast, disruption of genes PAS_chr4_0587 (SHM1) encoding mitochondrial (Shm1) did not reduce remarkedly the eGFP fluorescence (Fig S6). These findings substantiate the hypothesis that the intracellular formate is higher in the *fdh1*Δ strain, accounting for pAOX1 induction in non-repressive culture conditions.

**Figure 4:**
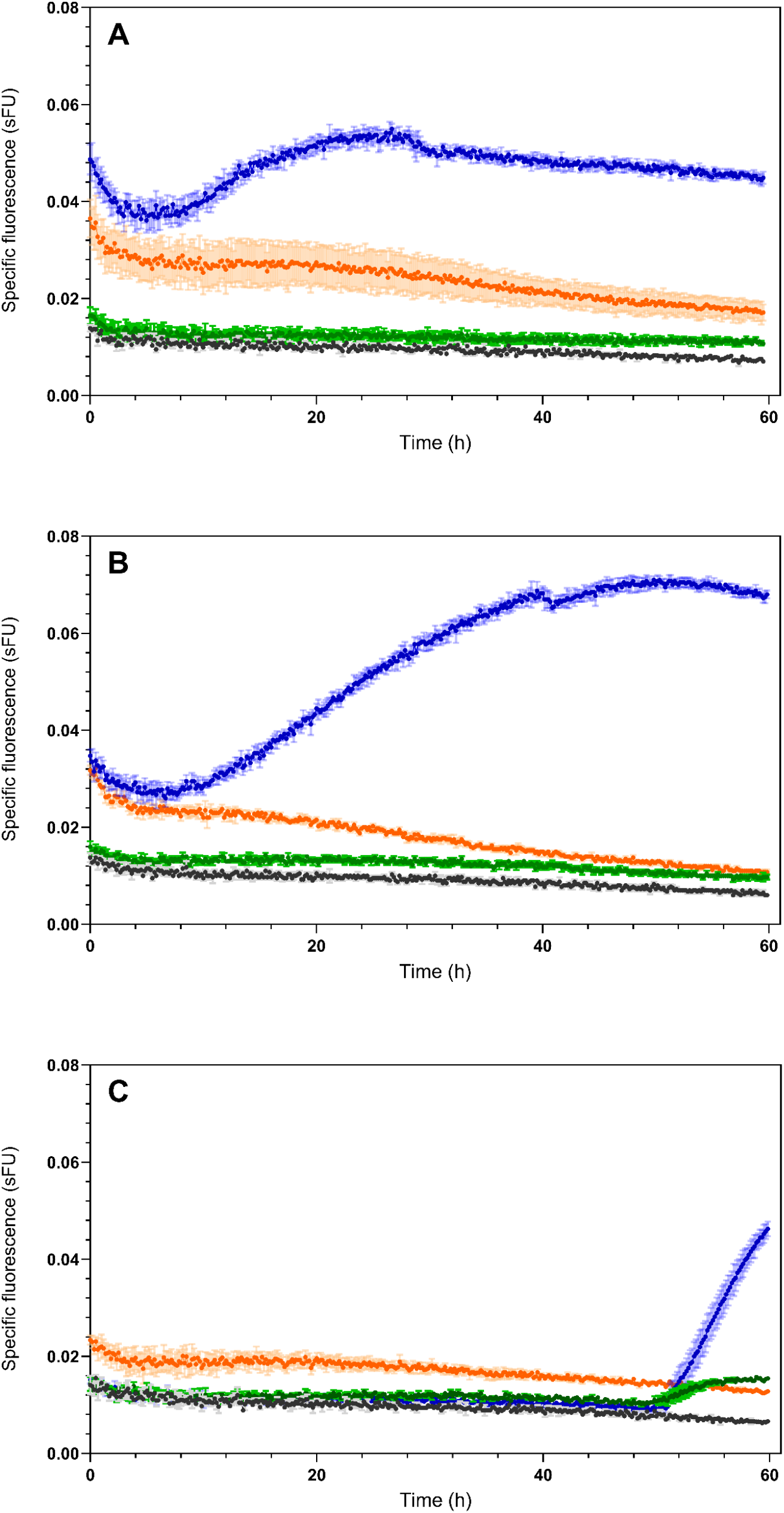
Specific eGFP fluorescence of RIY232 (WT, black); RIY230 (pAOX1-eGFP, green); RIY540 (fdhΔ, pAOX1-EGFP; blue); RIY640 (fdhΔ, shm2Δ pAOX1-eGFP; orange) strains; during growth in YNB minimal medium containing sorbitol (YNBS, panel A), sorbitol and serine (YNBSS, panel B), and sorbitol and glycine (YNBSG, panel C) Cells were grown in BioLector and specific fluorescence values are means and standard deviation on four cultures replicates. sFU: specific fluorescence unit.

### Production of a secreted protein by an *fdh*Δ mutant in sorbitol-based medium

In many recombinant protein (rProt) production processes using *P. pastoris*, glycerol is used in a first phase to generate biomass at a high cell density, to repress pAOX1 and thus to prevent rProt synthesis. In a second phase, the carbon source is shifted to methanol or to a mixture of methanol and sorbitol to trigger rProt synthesis by induction of pAOX1 promoter (Niu *et al*., 2013; Carly *et al*., 2016; Berrios *et al*., 2017). In the rProt production phase, the purpose is to direct most of the energy from carbon sources to rProt synthesis while minimizing cell growth. Herein, the lipase B from *Candida antarctica* (CalB) was used in combination with the α-mating factor from *S. cerevisiae* as a secretory protein reporter. The CalB coding sequence was cloned under the control of the pAOX1 promoter and integrated into the genome of the RIY232 strain, a prototroph derivative of *P. pastoris* GS115 (Velastegui *et al*., 2019). In the resulting RIY308 strain (*pAOX1-αMF-CalB*, CalB strain), the *FDH1* encoding gene was then knockout to yield the RIY561 strain (*fdh1Δ, pAOX1-αMF-CalB*, CalB-*fdh1Δ* strain). Both strains were grown either on glycerol, on a mixture of methanol and sorbitol (60/40, 0.3 C-mol as in Carly et al., 2016; Niu et al., 2013), and on sorbitol (i.e., YNBG, YNBMS and YNBS, respectively). Biomass and specific lipase CalB activity were quantified at different time points over 36 h (Fig.5).

On glycerol, cell growth for the CalB and the CalB-*fdh1Δ* strains were similar, with biomass values equal to 10.6 ± 0.1 and 10.1 ± 0.3 gDCW l^-1^, respectively, at the end of the growth phase (i.e., 12h, Fig 5.A). As expected, the lipase activity could not be detected during the first 12h, then after it increased slightly upon glycerol exhaustion in the medium (i.e. in pAOX1 derepressed condition, data not shown). On methanol (YNBSM), the biomass of the CalB-*fdh1Δ* strain was markedly lower compared to the CalB strain, most probably due to the accumulation of toxic methanol catabolism byproducts (i.e., formate) as previously reported (Guo *et al*., 2021). For both strains, the specific CalB lipase activity increased similarly over time to reach values after 30 h of 113.6 and 113.3 U mgDCW^-1^ for the CalB and the CalB-*fdh1Δ* strains, respectively (Fig 5.D). On sorbitol, both strains exhibited similar lower biomass values as compared to the glycerol medium. This could be lined with the lower uptake rate for sorbitol compared to glycerol (0.02 g gDCW^-1^ h^-1^ and 0.9 g gDCW^-1^ h^-1^, respectively; data not shown). Most importantly, the maximal specific lipase activity was remarkedly higher for the FDH disrupted strain (CalB-*fdh1Δ*) compared to the non-disrupted one (i.e. 130-fold). The specific lipase CalB activity for the CalB-*fdh1Δ* strain was in the same range on sorbitol and sorbitol-methanol medium (136 U mgDCW^-1^ and 113.3 U mgDCW^-1^, respectively). However, it was reached 2.4 times faster on sorbitol medium (i.e. after 15h and 36 h, respectively, Fig 5F).

**Figure 5.**
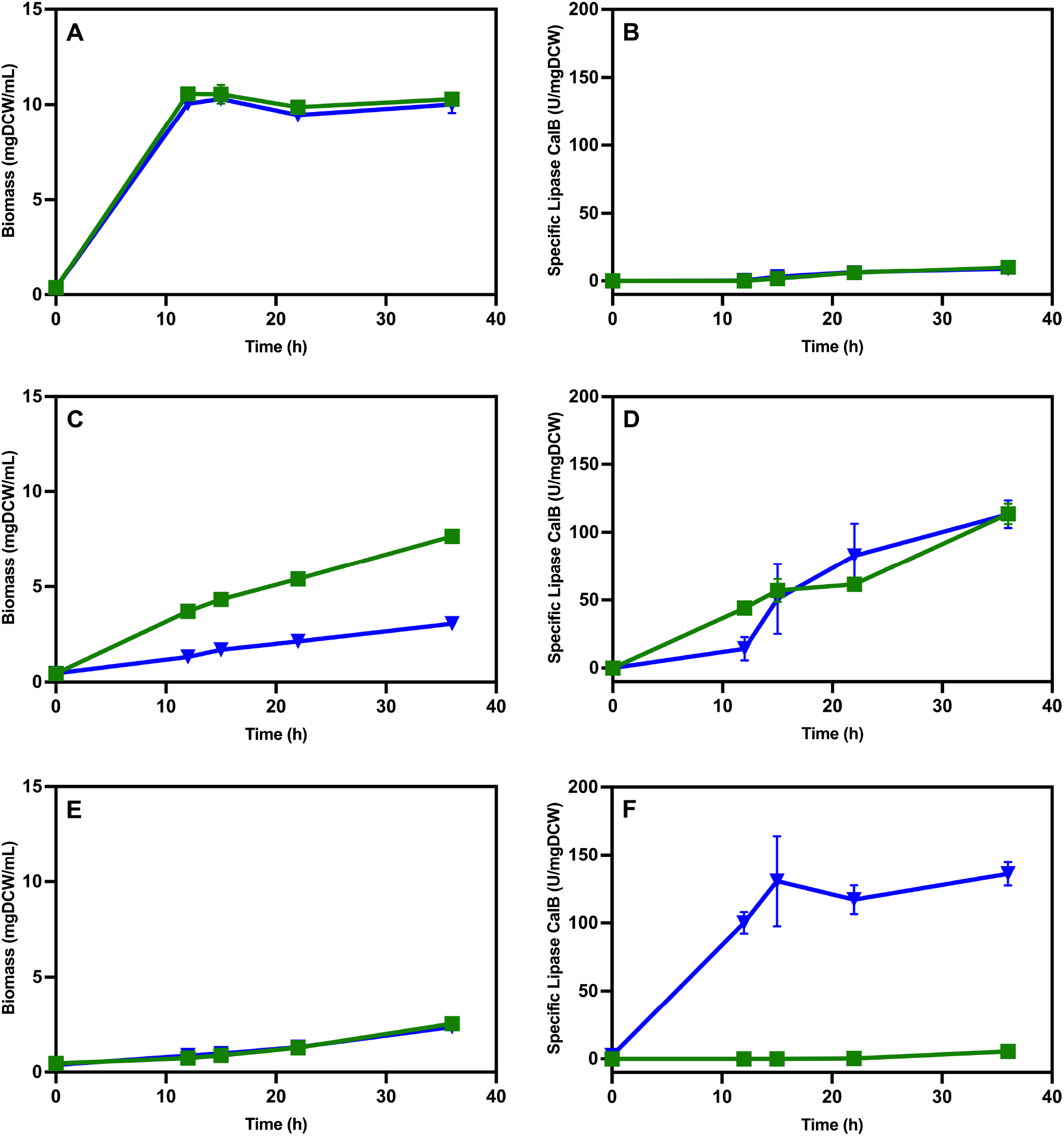
Biomass and specific lipase activity during growth of strains RIY 308 (pAOX1-αMF-CalB, strain CalB, green squares) and RIY561 (fdh1Δ-pAOX1-αMF-CalB, strain CalB-fdh1Δ, - blue triangles) in the presence of glycerol (YNBG, panels A and B), methanol-sorbitol (YNBMS, panels C and D) and sorbitol (YNBS, panels E and F). Data are mean and standard deviation from cultures were performed in triplicate in shake flasks in triplicate. Lipase assays were performed in triplicates.

## Conclusion

Herein, we have demonstrated that formate from the THF-C1 metabolism induces the pAOX1 promoter in an *fdh1Δ* strain grown under derepressed culture conditions. This is particularly interesting for recombinant protein production processes, as adding inducers such as methanol or formate is no longer required to trigger rProt synthesis. By growing the cells in a mixture of glycerol and sorbitol, rProt synthesis is initiated upon glycerol depletion in the medium. This autoinduced system paves the way for further development of methanol-free processes for rProt synthesis in *P. pastoris*.

## DATA AVAILABILITY

Data are available upon request to the corresponding author

## AUTHOR CONTRIBUTIONS

**Cristina Bustos:** Conceptualization; data curation; formal analysis; investigation; methodology; validation; visualization; writing – original draft; writing-review and editing. **Patrick Fickers:** Conceptualization; formal analysis; investigation; methodology validation; validation; visualization; funding acquisition; resources; supervision; writing – original draft; writing - review and editing. **Julio Berrios:** Conceptualization; funding acquisition; supervision; writing-review.

## ACKNOWLEDGEMENTS

The authors thank Prof. Gasser from CD Laboratory of growth-decoupled protein production in yeast, Department of Biotechnology, University of Natural Resources and Life Sciences for providing pKTAC-Cre vector. A. Anckaert, M. Delvenne, Vandenbroucke, V., S. Steels and R. Thomas are acknowledged for their technical help and fruitful discussion.

