## Supplementary figures and tables for "Formate from THF-C1 metabolism induces the AOX1 promoter in formate dehydrogenase-deficient *Pichia pastoris*"

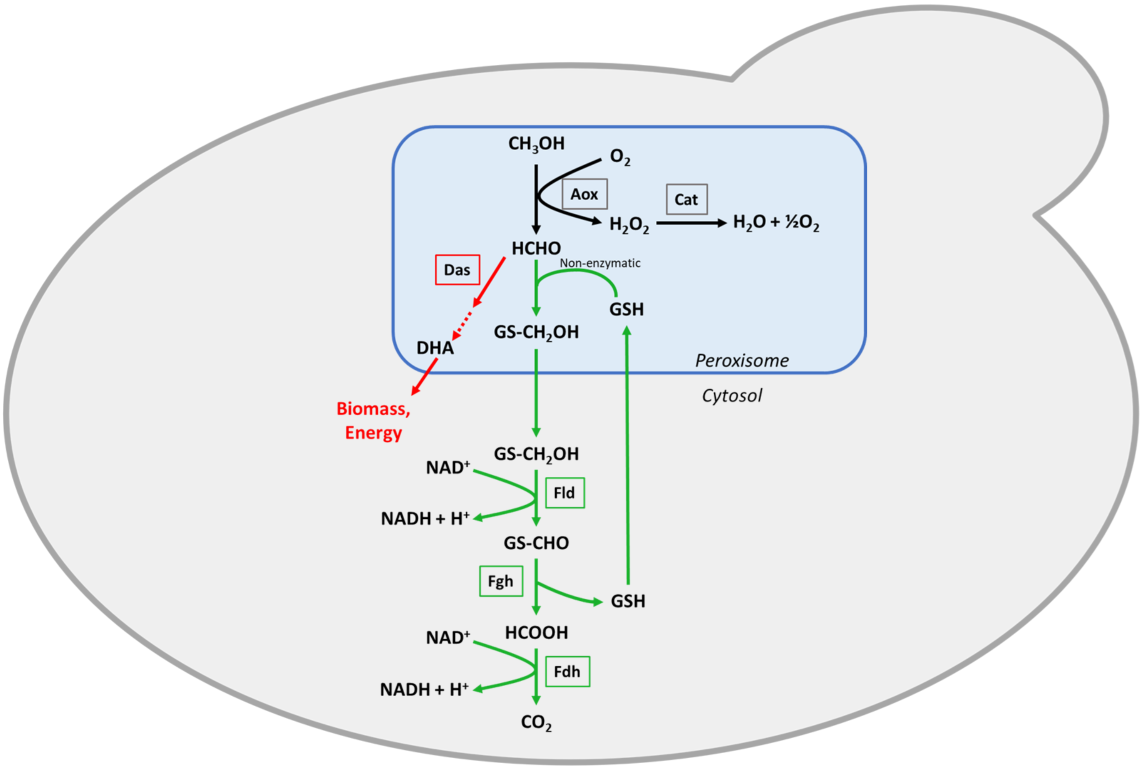


**Figure S1. Methanol pathway in *P. pastoris***. *Enzymes:* Aox: alcohol oxidase; Cat: catalase; Das: dihydroxyacetone synthase; Fld: formaldehyde dehydrogenase; Fgh: S-formylglutathione hydrolase; Fdh; formate dehydrogenase. *Abbreviations:* Gs-CH2OH: S-hydroxymethyl glutathione; Gs-CHO: S-formylglutathione; GSH: reduced form of glutathione. *Pathway*: Fld, Fgh and Fdh are parts of the methanol dissimilation pathway that convert formaldehyde (HCHO) into carbon dioxide ($CO_2$).

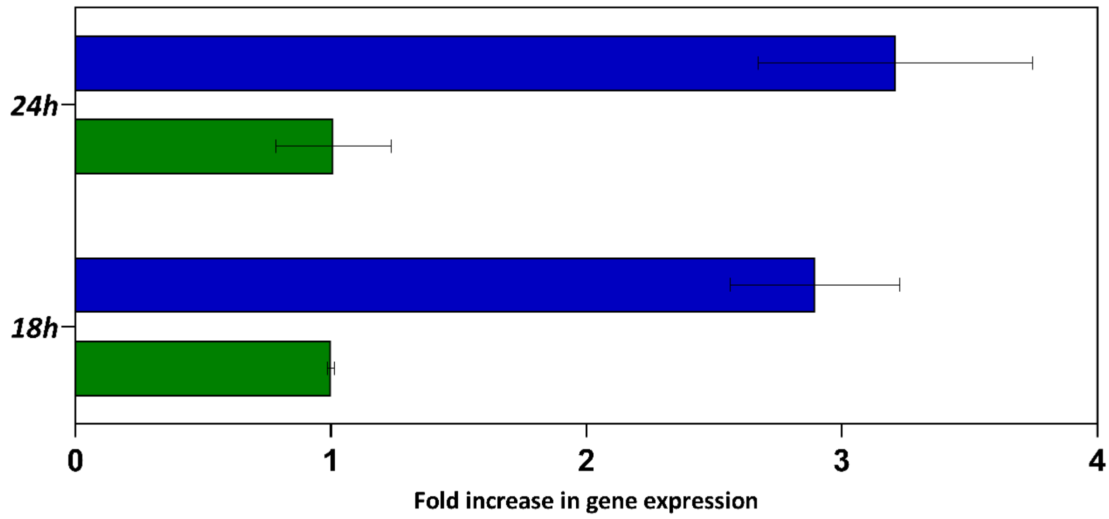


**Figure. S2.** *Relative expression level of eGFP gene in strain RIY230 (pAOX1-eGFP, strain FDH1, green) and RIY540 (fdh1Δ, pAOX1-eGFP, strain fdh1Δ, blue) in minimal medium containing sorbitol (YNBSC). Samples were collected after 18 h and 24h. Displayed values were normalized to that of the actin gene and corresponded to means and standard deviations from three independent replicates conducted in flake flasks.*

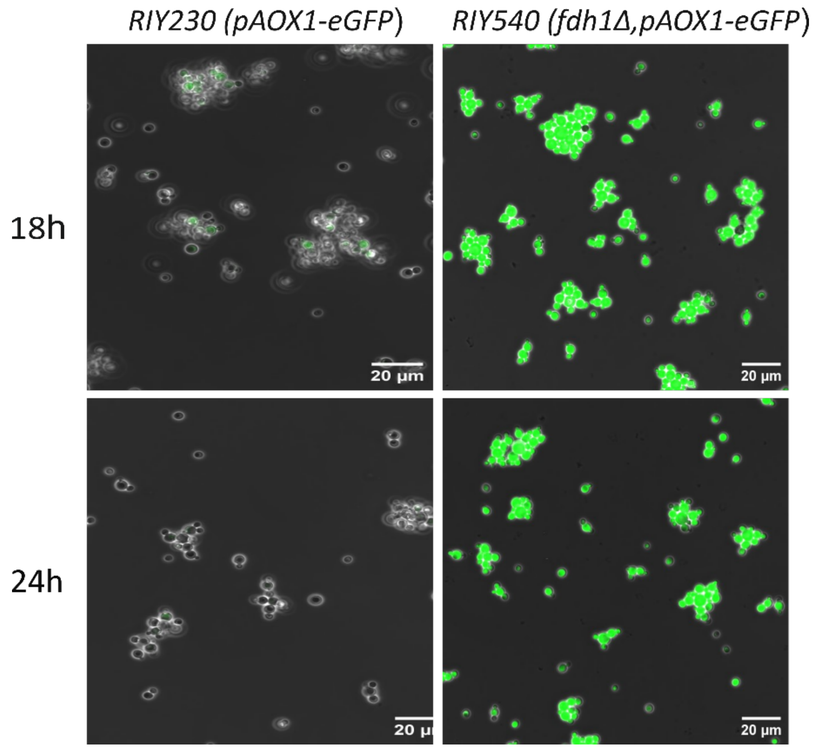


**Figure. S3.** *Observation of P. pastoris strains RIY230 (pAOX1-eGFP, strain FDH1) and RIY540 (fdhΔ, pAOX1-eGFP, strain fdh1Δ) after 18h and 24 h of growth in minimal medium containing sorbitol (YNBSC) by fluorescent microscopy. A representative sample from the triplicate cultures conducted in flake flask are shown. Observation and image processing are detailed in material and methods.*

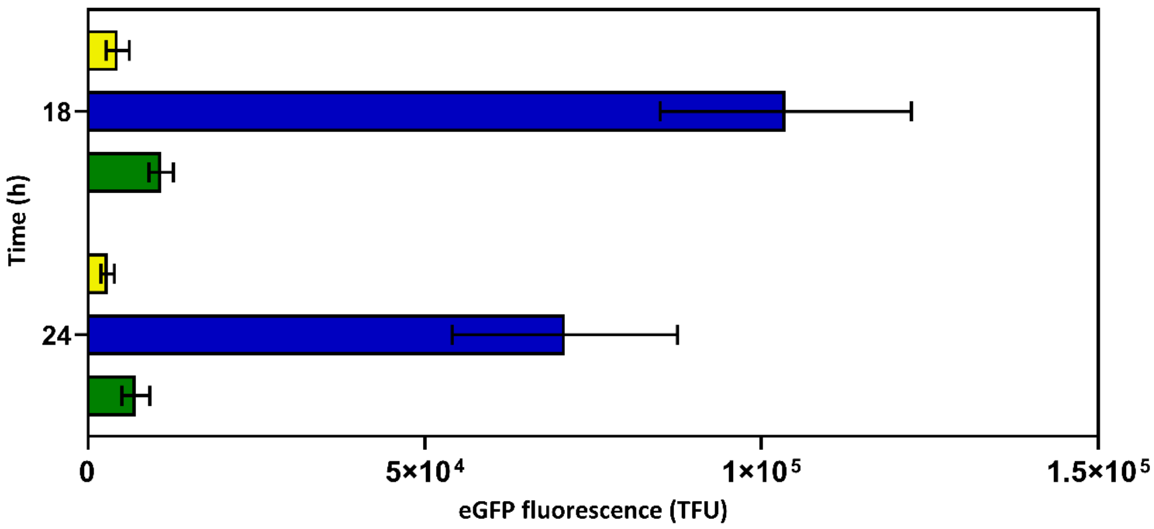


***Figure S4.*** *eGFP fluorescence of strain RIY230 (pAOX1-eGFP, strain FDH1, green); RIY540 (fdh1Δ, pAOX1-eGFP, strain fdh1Δ blue); RIY624 (fdh1Δ, pAOX1-eGFP, pGAP-FDH, strain, fdh1Δ-FDH1, yellow) after 18h and 24 h of growth in minimal medium containing sorbitol (YNBSC). Fluorescence was quantified by flow cytometry on 20,000 cells and expressed as TFU (total fluorescence, see materials and method for calculation details). Values are the means and standard deviation from biological triplicate cultures conducted in shake flasks.*

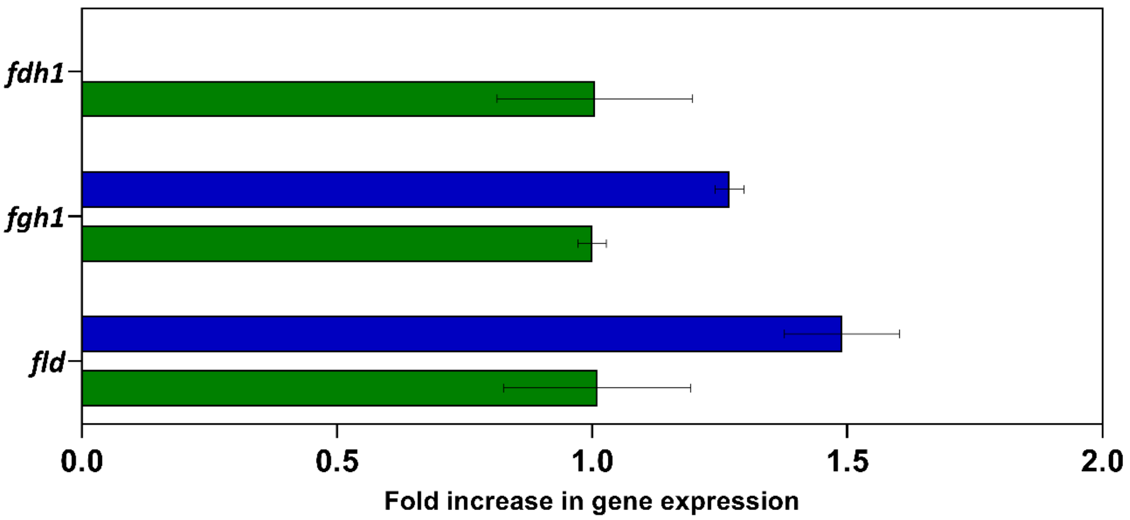


***Figure S5.*** *Relative expression level of genes involves in the methanol dissimilation pathway FLD (PAS_chr3_1028), FGH1 (PAS_chr3_0867), FDH1 (PAS_chr3_0932) in the strains RIY230 (pAOX1-eGFP, strain FDH1, green) and RIY540 (fdhΔ1, pAOX1-eGFP, strain fdhΔ1, blue) in YNBSC medium. Samples were collected after 18 h of culture. Displayed values were normalized to that of the actin gene and corresponded to means and standard deviations from duplicates independent replicates cultures conducted in flake flasks.*

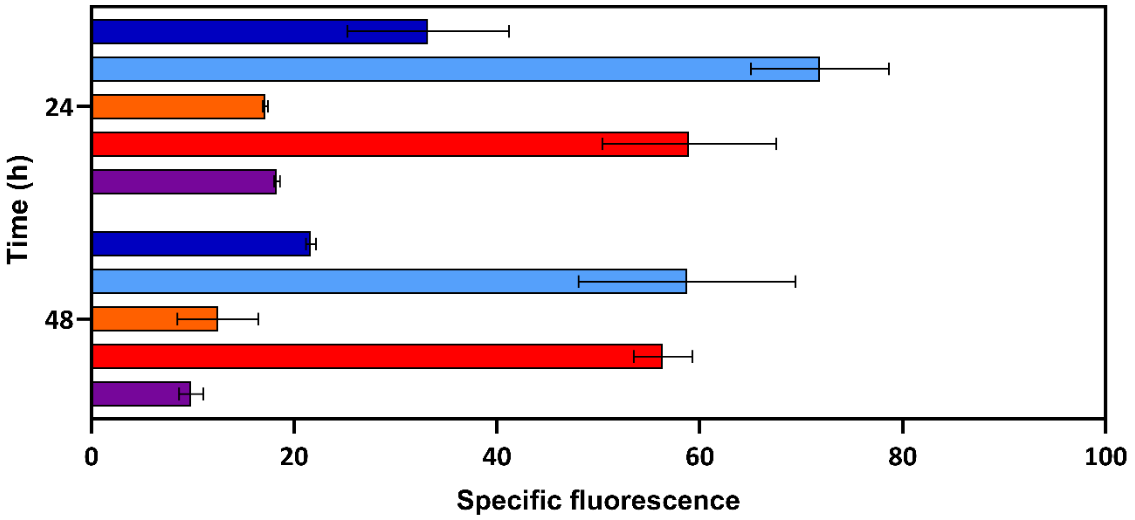


**Figure. S6. Specific eGFP fluorescence of strain** *RIY540 (fdh1Δ, pAOX1-EGFP; growth in YNBS, blue); RIY540 (fdh1Δ,, pAOX1-EGFP; growth in YNBSS, light blue); RIY640 (fdh1Δ, shm2Δ pAOX1-eGFP; growth in YNBSS, orange); RIY641 (fdh1Δ, shm1Δ, pAOX1-eGFP, pGAP-FDH; growth in YNBSS, red); RIY642 (fdh1Δ, shm1Δ ,shm2Δ ,pAOX1-eGFP, pGAP-FDH; growth in YNBSS, purple). Values are the means and standard deviation from two biological replicates conducted in shake flasks. Biomass measures and specific fluoresce unites quantification was detailed in materials and methods.*

Table S1. *Escherichia coli* strains used in this study.

| Name | Plasmid - genotype | Source/Reference |
|---|---|---|
| A2 | BB1_23 | (Prielhofer *et al.*, 2017) |
| D12 | BB3aZ_14 | (Prielhofer et al., 2017) |
| A4 | BB1_12_pGAP | (Prielhofer et al., 2017) |
| C1 | BB1_34_ScCYC1tt | (Prielhofer et al., 2017) |
| E1 | BB3eH_14 | (Prielhofer et al., 2017) |
| E6 | BB3aN_14 | (Prielhofer et al., 2017) |
| RIE396 | pKTAC-Cre | (Marx et al., 2008) |
| RIE369 | RIP369, pGEMTeasy, *FDH1* disruption cassette | This work |
| RIE465 | RIP465, BB1-23-*FDH1* | This work |
| RIE466 | RIP466, BB3eH_14, *pGAP-FDH-scCYC1tt* | This work |
| RIE 491 | RIE491, TopoBluntII, *SHM2* disruption cassette, Zeo | This work |
| RIE492 | RIP492, TopoBluntII, *SHM1* disruption cassette, Nat | This work |

Table S2. Primers used in this study

| Name | Sequence 5´to 3´ | Restriction site |
|---|---|---|
| M13-Fw | GTAAAACGACGGCCAGT | |
| M13-RV | AACAGCTATGACCATG | |
| P.fdh1-Fw | GGGCAGAAGGATCAGCCTGGACGAAG | |
| P.fdh1-Rv | GGGGA**GGTCTC**ACCTGCGTGTTTAAGTGGGTGATGT | BsaI |
| BleoR.fdh1-Fw | GGG**GGTCTC**GCAGGTCGACAACCCTTA | BsaI |
| BleoR.fdh1-Rv | GGGC**GGTCTC**ACTTCAGTGACAACGTTGCTGAAGCAGT | BsaI |
| T.fdh1-Fw | GGC**GGTCTC**TGAAGTGACTTTATGAATTCGCAA | BsaI |
| T.fdh1-Rv | GGGGTAGCCTCAACAATTGGCAGCTCTTC | |
| Up.fdh1-Fw | AGAAGAGCATCTCAACTATGCCTATG | |
| BleoR.Int-Rv | CATGGTTTAGTTCCTCACCTTGTC | |
| BleoR.Int-Fw | GGAGCAGGACTGATCAGTACTTACTGA | |
| Dw.fdh1-Rv | GTTCAATGACGAAAAGGTGGTGTTGG | |
| Fdh1-Fw | AACC**GGTCTC**ACATGAAAATCGTTCTCGTT | BsaI |
| Fdh1-Rv | ACC**GGTCTC**CAAGCTTTATGCGACCTTTTTG | BsaI |
| Fdh1.BpiI-Fw | TACTACGACTACCAAGGTCTGCCAAAAGAG | |
| Fdh1.BpiI-Rv | CTCTTTTGGCAGACCTTGGTAGTCGTAGTA | |
| pGAp.Int-Fw | CGTCGCTGGCAATAATAGCGG | |
| Cyc1t.Int-Rv | GGGACCTAGACTTCAGGTTGTC | |
| P.shm1-Fw | GCATTCCGGAAATAAATCATATGT | |
| P.shm1-Rv | GCGTCTTCCTTGTTGTGCTTTTCTTTCAATAGTAGAG | |
| Nat.shm1-Fw | AGCACAACAAGGAAGACGCCGCTCC | |
| Nat.shm1-Rv | CTATAGTTTAATTGTTTTCAGTGACAACGTTGCTGA | |
| T.shm1-Fw | AACGTTGTCACTGAAAACAATTAAACTATAGGTGCCTTACT | |
| T.shm1-Rv | CCTCATCACTGAACAATCTGAG | |
| Up.shm1-Fw | GCATTGGAAAAGATCGTTTTTATTTG | |
| Dw.shm1-Rv | GGTATTTGCATGATAGTTTTATCCATTTC | |
| Nat.Int-Fw | CTGACCAAGGTGTTCCCC | |
| P.shm2-Fw | TGCAACCTGAGATCTTGAGACA | |

| | | |
|---|---|---|
| P.shm2-Rv | AGGGTTGTCGACCTTTATTTGGATAGGTGGGTAGTTTGG | |
| BleoR.shm2-Fw | CACCTATCCAAATAAAGGTCGACAACCCTTAATATAAC | |
| BleoR.shm2-Rv | TCACTAATTATATTCGTGGATCTGATATCACCTAATAAC | |
| T.shm2-Fw | TGATATCAGATCCACGAATATAATTAGTGAACAAAAGAATATAAA TAA | |
| T.shm2-Rv | GTAATTTCTGCTTCCGGTTCTT | |
| Up.shm2-Fw | CAAGGTTAACGGTTCACCTATC | |
| Dw.shm2-Rv | TTCAAATCTTCCAACCCAACTTC | |
| qAct-F | AGATGGCTCCGAGAAGTTCA | Actin |
| qAct-R | GTTGCTCAGAGGGCTTCAAC | Actin |
| qFLD-F | ATCACTGACGGAGGCTTTGA | FLD |
| qFLD-R | TGGCATTTGAGTACGTCCCT | FLD |
| qFGH-F | CCCAAATTGCAGGCTGACTT | FGH |
| qFGH-R | AGTGGGGTTGGAGATTGGAG | FGH |
| qFDH-F | GCCGATGTTGTTACCGTCAA | FDH |
| qFDH-R | GTCACCACCGTAACCTCTCA | FDH |
| qeGFP-F | ATCATGGCCGACAAGCAGAA | EGFP |
| qeGFP-R | TCTCGTTGGGGTCTTTGCTC | EGFP |
